# Size-dependent sex allocation and the expression of andromonoecy in a protogynous perennial herb: both size and timing matter

**DOI:** 10.1101/2023.03.10.532080

**Authors:** Kai-Hsiu Chen, John R Pannell

## Abstract

The optimal life history and sex allocation of perennial hermaphrodites should depend on both their size and the relative costs and benefits of reproducing through male versus female functions. Theory predicts that insect-pollinated perennials should increase their allocation to female function with size, while the ‘mating environment’ hypothesis predicts that allocation to male function should track mating opportunities over the course of flowering. We test these two predictions by inferring male and female reproductive success in the protogynous perennial herb *Pulsatilla alpina* by tracking the patterns and dynamics of sex allocation over time for marked individuals over a range of sizes. We found that small individuals tend to produce only male flowers and that both small and larger individuals produced male flowers at the beginning of the flowering season when mating opportunities were high. By considering within-population variation in life history and phenology jointly rather than separately, and by considering both the tradeoff costs and benefits of allocation to male versus female functions, our results provide new insights into the evolution of both gender diphasy and andromonoecy in perennial plants that are constrained by a dichogamous flowering strategy.

## Introduction

Perennial hermaphroditic plants face a complex set of allocation decisions that affect their life history and sex allocation – the two principal axes of reproduction. First, they must determine when, and at what size, to begin to flower, and whether then to invest in reproduction more heavily in some years over others (aspects of their life history) (Lovett Doust 1989; Wenk and Falster 2015; Roach and Smith 2020). Second, they must determine how much of their reproductive resources to allocate to their male versus female sexual functions (their sex allocation, ‘SA’) (Charnov 1982; Charlesworth and Morgan 1991). Life-history and sex-allocation decisions are each important on their own, but they should also be linked because tradeoffs between growth and reproduction will often differ through reproduction through male versus female functions (Iwasa 1991; Obeso 2002; Dorken and Van Drunen 2018). Empirical assessment of the marginal costs of male versus female function in terms of growth and survivorship indicates that plants often invest more heavily in their female function than their male function (Obeso 2002). This is particularly evident in dioecious species in which adult sex ratios of reproducing individuals are male-biased as a result of the greater mortality or less frequent flowering of females that bear a higher cost of reproduction (Sinclair et al. 2012; Barrett and Hough 2013; Field et al. 2013), but it is also evident in hermaphroditic species in which plants that have invested heavily in their female function are less likely to flower the following year than those that have reproduced as males (Schlessman 1991; Zhang et al. 2014; Blake-Mahmud and Struwe 2019; Bialic-Murphy et al. 2020).

Whereas patterns of SA in plants are often interpreted from the perspective of the relative costs of male versus female functions (Goldman and Willson 1986; Case and Ashman 2005), selection optimizes phenotypes in terms of the costs *and benefits* of the strategies they adopt. Thus, while a greater cost of female function has been invoked to explain patterns of size-dependent SA in which plants emphasize their male function when small and their (costlier) female function when large (Lloyd 1984), these patterns must also depend on the relative benefits to small versus large individuals of reproducing through male versus female function (Ghiselin 1969; Klinkhamer et al. 1997). In insect-pollinated plants, fitness gained through allocating to male function is thought to saturate more quickly than through female allocation, so plants with a large budget should have a female-based SA (de Jong and Klinkhamer 1989, 1994). In contrast, tall plants of wind-pollinated species may enjoy direct effects of fitness because height is expected to enhance pollen dispersal, so we might expect large size to be associated with the male function (de Jong and Klinkhamer 1994; Sakai and Sakai 2003; Cadet et al. 2004). Importantly, in species with wind-dispersed fruits or seeds, female fitness might also benefit from the direct effects of height (de Jong and Klinkhamer 1994; Soons et al. 2004; Pickup and Barrett 2012).

The relative benefits of flowering through male versus female functions may also vary over time *within* a flowering season (Brunet and Charlesworth 1995). Because all flowers in a population do not open simultaneously, the pollen from flowers produced at different times during the flowering season will have different opportunities to sire offspring, and the intensity of male-male competition will vary accordingly; we might refer to this idea as the ‘mating environment’ hypothesis (Brunet and Charlesworth 1995). In protandrous species, where the male function precedes the female function, more females will be available late in the season, and we should therefore expect some plants to capitalize on the greater availability of female mates by emphasizing their male function towards the end of the flowering season. An increase in relative allocation to male function has indeed been reported for some protandrous species (but see Aizen 2001), e.g., *Aquilegia caerulea* (Brunet 1996), *Campanula rapunculoides* (Vogler et al. 1999), *Cimicifuga simplex* (Pellmyr 1987), and *Corydalis ambigua* (Kudo et al. 2001). In protogynous species, by contrast, the mating environment hypothesis predicts an increased allocation of resources to male function by some individuals early in the season (Brunet and Charlesworth 1995).

While the mating environment hypothesis provides a plausible explanation for an increase in relative allocation to male function in protandrous species late in the flowering season, alternative theories predict the same pattern. For instance, the ‘resource competition’ hypothesis proposes that flowers later in a season have fewer resources to draw upon due to the allocation of resources into fruiting by early-season flowers and that late flowers should avoid the costly female function and emphasize their male function (Diggle 1994; Medrano et al. 2000). Alternatively, the ‘architecture effect’ hypothesis suggests that resources are more available to flowers at the base of the inflorescence, which are usually the early flowers (Wyatt 1982; Diggle 1995), such that an acropetal decline in allocation towards female function along the inflorescence should be expected. Significantly, although these hypotheses provide alternative explanations for within-season shifts in allocation in protandrous species, only the mating environment hypothesis predicts an increase in relative allocation to male function in early flowers of protogynous species at both the individual and flower levels (Brunet and Charlesworth 1995; Huang et al. 2004). As a consequence, protogynous species provide better systems to test the mating environment hypothesis of SA.

The predicted higher relative allocation to male function in early-season flowers of protogynous species has rarely been tested (Austen et al. 2015). A decline in anther number in later flowers within individuals has been found in protogynous *Aquilegia yabeana* (Huang et al. 2004) and *Helleborus foetidus* (Guitián 2006), but these studies did not assess the temporal dynamics of mate availability and prospective male reproductive success (‘RS’) over the course of a flowering season (though see Gleiser et al. 2008). We also know little about how SA varies as a function of *both* size and phenology jointly. A few empirical studies have reported an extreme pattern of shifting from functionally a pure male to a hermaphroditic strategy with size (Schlessman 1991; Kudo and Maeda 1998; Peruzzi et al. 2012; Zhang et al. 2014; Niu et al. 2017), but we remain largely ignorant about the fitness of the functionally pure males. We are particularly ignorant of how plants respond to changing opportunities of mating within seasons, and of how the strategy they adopt depends on their size.

In this study, we ask how perennial plants respond to the changing costs and benefits of allocation to male versus female sexual functions over time between and within seasons. Specifically, we studied the size, flowering behavior, and SA of marked individuals of the strongly protogynous alpine herb *Pulsatilla alpina* (Ranunculaceae) over consecutive seasons, as well as in detail over time within a reproductive season, and we used our observations to evaluate the mating environment hypothesis on the basis of inferred male and female fitness components. We were particularly interested in seeking an explanation for (1) the expression of gender diphasy in *P. alpina*, in which small (and likely young) individuals produce only male flowers and larger (older) individuals are hermaphroditic, and (2) the evolution of the heteromorphic sexual system as andromonoecy, where plants produce purely male flowers in the context of an overall hermaphroditic SA strategy. Our study illustrates the value of a dynamic assessment of both life history and SA jointly in terms of the changing costs and benefits of mating as male versus female.

## Materials and methods

### Study species and study sites

*Pulsatilla alpina* (L.) Delarbre (Ranunculaceae) is a perennial, protogynous, andromonoecious hemicryptophyte that grows in sub-alpine to alpine habitats in central Europe (Lauber et al. 2018). Several vegetative and/or reproductive shoots emerge from a perennial underground rhizome soon after the snowmelt, from early May to July. Individuals produce up to approximately twenty white flowers, each on its own reproductive shoot. Phenotypically male flowers bear only stamens, whereas protogynous hermaphroditic flowers bear stamens and uni-ovulate pistils. In the populations studied here, stamen and pistil numbers varied between approximately 150 and 400, and zero and 400, respectively. The flowers are predominantly visited by dipteran insects, including houseflies and syrphid flies (Chen and Pannell 2022). Ripe achenes with elongated pappus hair are dispersed by wind in early autumn (Vittoz and Engler 2007). After the achenes are dispersed in late autumn, the above-ground vegetative parts of individuals die away, but individuals persist as rhizomes below ground until the next spring.

We studied key aspects of the life history, SA, and reproductive biology of *P. alpina* in several populations in the pre-Alps of the canton Vaud, Switzerland during the flowering seasons of 2018 to 2020: one population at Les Mosses (Population LM), two populations at Lac Lioson (Populations LL1 and LL4), and two populations at Solalex (Populations S1 and S2). Population LM (625 individuals) is located on an open slope of sub-alpine grassland surrounded by forest, covering an area with dimensions of about 200 m x 50 m. Populations LL1, LL4, S1, and S2 (all > 1,000 individuals) are located on open slopes of sub-alpine grassland (see Appendix table S1 for a more detailed description of the populations). In Population S1, we tracked flowering behavior and the SA of individuals over two consecutive seasons to investigate how individuals shift their SA. In Population LL1, we assessed the siring ability of pollen from male and hermaphrodite flowers (outcross and self-pollination). In Population LM, we tracked the flowering status, SA, and mating opportunities of marked individuals over the course of the flowering season to relate components of RS to plant size and the timing of reproduction through male and female functions. Finally, we measured SA and mate availability again in 2019 in Populations LL1, LL4, S1, and S2 to determine the extent to which patterns found in Population LM are similar among different populations with different size distributions.

### Siring ability of pollen from different sources

In Population LL1 we conducted hand pollination treatments in 2018 to assess the ability of pollen from male and hermaphrodite flowers of *P. alpina* to sire progeny and to compare siring ability via outcrossing versus selfing. Individuals with more than one floral shoot were chosen at the beginning of the flowering season. Floral buds of hermaphrodite flowers were bagged with tea bags as soon as the floral shoots emerged until opening. Anthers in the flower were removed to avoid self-pollination. The pollination manipulations were conducted after the flower opened and before the anthers dehiscence, that is at the female stage (see Appendix table S2 for details). The pistils in the flowers were hand-pollination with pollen from a flower from the same individual (selfing, *N* = 10), from a male flower in another individual (outcrossing, *N* = 10), or from a hermaphroditic flower in another individual (outcrossing, *N* = 12). The pollen used in outcrossing manipulations was from individuals at least 5m away from the hand-pollinated individuals. After the manipulation, the flowers were bagged again until fruiting. In addition, open-pollinated flowers were labeled as control (*N* = 15). In early autumn, all parts of the seed head were collected from the plants before dispersal. The seed set was counted as the mature seed number divided by the total pistil number.

### Phase changes in individual SA from one season to the next

To assess how the SA of individuals changes among seasons, we marked 111 individuals in Population S1in 2019 (as described above for Population LM), including in our sample both flowering and non-flowering individuals and the full range of sizes and SA. The SA at the flower and the individual levels was quantified in 2019 and 2020, as described below for Population LM.

### Flowering phenology

In Population LM, we studied flowering phenology in 2018 by recording the flowering state of all 625 individuals every three or four days throughout the flowering season, from early May to late June. All flowering individuals in the population were marked with a metal tag nailed into the ground beside each plant. Flowers were individually labelled with a paper tag. For each flower, we recorded its sexual stage (female or male) at each time point from its opening until it wilted. Furthermore, we recorded the SA of each flower using four ordinal categories based on visual inspection of the pistil number, i.e., zero, one to 50, 51 to 150, and more than 150 pistils. For a subsample of 88 individuals, we also photographed each flower at the late female or early male stage to count the number of stamens and pistils (see details below). The flowering date for each flower was calculated as the mid-date between its opening and its wilting, while the flowering date for each plant was calculated as the mid-date between the opening of its first flower and the wilting of its last flower.

### Plant size

In Population LM, we estimated aboveground biomass (hereafter, size) at the end of the flowering season in 2018 by harvesting all aboveground parts of a subsample of 88 of the 625 individuals in the population. The subsampled tissues were then dried in an oven for five days at 60LJ and then weighed. Individuals produce new aboveground tissue each growing season from resources stored in the rhizome and root tissues. Although belowground biomass was not estimated in this study (it would have required killing the plants), belowground and aboveground biomass is typically correlated in plants (Müller et al. 2000; Enquist and Niklas 2002).

### Estimates of SA and functional gender

In Population LM, we further counted the number of stamens and pistils produced in each flower, estimated the absolute biomass allocated to each sex and used these data to calculate the functional gender for each of the 88 individuals sampled (Lloyd 1980; see below). To estimate female allocation, we collected and counted all pollinated and unpollinated pistils in each flower at the end of flowering from July to August (see estimates of seasonal female RS below). To estimate male allocation, we counted stamens photographed on each flower at the time of harvest. To account for potential biases in counting floral parts on the basis of photographs rather than the samples themselves, we counted the number of stamens directly on 15 flowers and determined the relation between the two counting methods using regression, which was: *m_R_* = 1.66*m_P_* + 27.2 (*r*^2^ = 0.654), where *m_R_* and *m_P_* are the estimates of stamen number on the basis of direct and image-based counts, respectively. We calculated the absolute allocation to male and female functions by multiplying the number of pistils and stamens by the mean weight of one pistil (4.41 x 10^-4^ g, estimated by averaging over 108 pistils) and one stamen (6.82 x 10^-5^ g, estimated by averaging over 770 stamens).

We estimated the functional gender (femaleness) at the individual level for all 625 individuals in Population LM based on the ordinal allocation categories mentioned below, and for all 88 sampled individuals based on actual counts of pistils and stamens. For both these estimates, we calculated individual functional gender according to Lloyd (1980) as

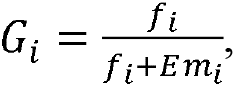

where *f_i_* is the number of pistils, *m_i_* is the number of stamens produced by the *i*th individual, and *E* = ∑ *f_i_* /∑ *m_i_* is an equivalence factor that accounts for the fact that the total number of genes transmitted through male and female functions must be equal at the population level. For the full sample of 625 individuals whose allocations were estimated in terms of ordinal allocation categories of the flowers produced, we determined *f_i_* as 0, 25, 100, and 225 pistils for the four ordinal categories, respectively, and *m_i_* as 204 stamens for all of the four sex-allocation categories, as we did not find any difference in the stamen number between male and hermaphroditic flowers. We then used these data to characterize the gross distribution of gender throughout the population. In addition, we used the more detailed data from the 88 subsampled individuals to relate the gender of plants to their size and flowering time during the season.

### Estimates of seasonal female RS

To estimate the seasonal RS via female function in Population LM (hereafter, female RS), we collected the achenes produced in each flower at the end of the growing season, using the same sample of 88 individuals measured for size and SA described above. We separated achenes into immature and mature categories and weighed. The immature achenes included non-fertilized pistils and achenes that had been damaged (killed) by seed predators (for details, see Chen and Pannell, 2022). We then estimated the female RS as the number of mature achenes produced.

### Estimate of seasonal male RS

To infer the prospective seasonal male RS in Population LM, we assumed a mass-action model of mating within a series of time windows over the course of the flowering season, which is commonly used to estimate the male RS in large wild populations in which estimates based on genotyping is unrealistic (e.g., Policansky 1981; Kudo and Maeda 1998). At each of the successive time points sampled, we assumed that each individual in the population could expect to sire an equitable fraction of all the ovules available to be fertilized within that time window, with the fraction calculated as the number of mature stamens on the individual at that point in time divided by the total number of stamens across the population. We calculated prospective male RS by multiplying this fraction with the number of pistils available at that time and with the mean mature seed set of the population, which is a constant independent of time (Appendix figure S1). Our estimate assumed no selfing at the flower level (autogamy is likely to be minimal because the species is strongly protogynous), but we did allow for geitonogamous selfing if the male and female stages of two flowers from the same individual overlapped (at a rate determined by random mating, according to the same the mass action model assumed for all other flowers). To calculate the total prospective male RS for each individual over the flowering season, we then summed the siring success inferred for each time window over the entire flowering for that individual. We ensured that the total male and female components of RS summed over all individuals over the whole flowering season were equal, as must be the case (given that all seeds have both a mother and a father).

### Patterns of mate availability and SA among populations

To determine whether the observed patterns of pistil availability and SA in Population LM (in 2018) reflect those in other nearby populations, we monitored flowering phenology, estimated SA, and measured plant size with the same methods described above in Populations LL1, LL4, S1, and S2 in 2019; these populations were larger and had a wider range of plant sizes than Population LM. In each population, we sampled around 70 flowering individuals of different sizes and labeled around 30 non-flowering individuals. The phenology of the populations and SA of the individuals were recorded once or twice a week through the flowering season, as described above. All aboveground tissue of the sampled individuals was harvested at the end of the growing season for weighing and analysis.

### Statistical analysis

All statistical analysis was conducted in R version 4.0.3 (R Core Team 2021). A generalized linear mixed model (*glmer* function in the R package *lme4* R; Bates et al., 2015) was built to evaluate the effects of pollination treatments on mature seed set. The mature seed set of each flower was set as a response variable with a binomial error distribution. Hand-pollination treatment (with four levels) was set as the fixed effect. Flower identity was set as a random effect to account for the non-independence of seeds from the same flower (the experimental unit was one seed). We used a posthoc Tukey test (*glht* function in the R package *multcomp*; Hothorn et al., 2008) to test the difference in mature seed set among treatments.

A generalized least square model (*gls* function in *nlme* package; Pinheiro et al., 2022) was built to evaluate the effects of size and flowering date on the absolute SA between female and male sex functions at the individual level, with absolute SA set as a response variable. In all analyses, size was transformed as log(size + 1) and then standardized to a mean of zero and a standard deviation of one. The flowering date was also standardized to a mean of zero and a standard deviation of one. Standardized size, standardized flowering date, and sexual function were set as the fixed effects, with three two-way and one three-way interaction terms. Variance in the absolute SA was allowed to differ between male and female functions. Individual identity was set as a random effect to account for the non-independence of absolute allocation to female and male functions within the same individual.

To evaluate the effects of size and flowering date on functional gender at the individual level, we built a linear model using the *lm* function in R, with functional gender set as the response variable and standardized size and flowering date together and their interaction set as fixed effects. To evaluate the effects of size and flowering date on female and male RS at the individual level, a *gls* model was used with the same model structure and relative RS set as a response variable. The relative female and male RS for each individual were calculated by dividing our estimates of the female or male RS, respectively, of each given individual by the corresponding mean RS across the sampled individuals.

We used a *gls* model to access the effects of size and flowering date on the seasonal total RS of different gender, i.e., male-phase and hermaphrodite-phase individuals. Relative total RS was set as a response variable, calculated by dividing the sum of female and male RS of an individual by the mean of the sum of female and male RS across the sampled individuals. Gender, standardized size, and standardized flowering date were set as fixed effects, along with their two-way and three-way interactions. The variance of relative total RS was allowed to differ between male-phase and hermaphrodite-phase individuals.

To test whether the earliest flowers produced by an individual had a higher probability of being entirely male, we used a generalized linear mixed model, with the probability of being a male flower set as a response variable. The flowering order of each flower from individuals producing a mixture of male and hermaphroditic flowers in Population LM was calculated according to the flowering date of the flowers. Individual identity was set as a random effect. We used a posthoc Tukey test to test the difference in male flower probability among flowering orders.

## Results

### Siring ability of pollen from different sources

We assessed the self-compatibility and relative siring ability of pollen from male versus hermaphroditic flowers in Population LL1. Flowers pollinated with pollen from both the same and from different individuals all produced seeds and fruit, confirming that *P. alpina* is self-compatible. There was no difference in siring ability between the pollen from male and hermaphroditic flowers (Figure S2). Outcrossed flowers had a significantly higher mature seed-set than selfed flowers (Figure S2), pointing to likely inbreeding depression at the seed development stage, which we calculated as 0.15. Open-pollinated flowers had a significantly lower mature seed set compared to hand-pollinated flowers (Figure S2), indicating a certain degree of pollen limitation.

### Phase changes in the SA of individuals from one season to the next

A total of 111 individuals were marked in 2019 in Population S1, among which 22, 32, and 57 individuals were in the non-flowering phase, male-phase, and hermaphrodite-phase, respectively. All 111 individuals were still alive in 2020. A total of 13 of them (12%) changed between male-phase and hermaphrodite-phase between years (Figure 1). A total of 4 (13%) and 46 (81%), respectively, of male-phase and hermaphrodite-phase individuals, remained in the same phase. See Table S3 for the transition matrix of marked individuals in different phases between two years.

**Figure 1.**
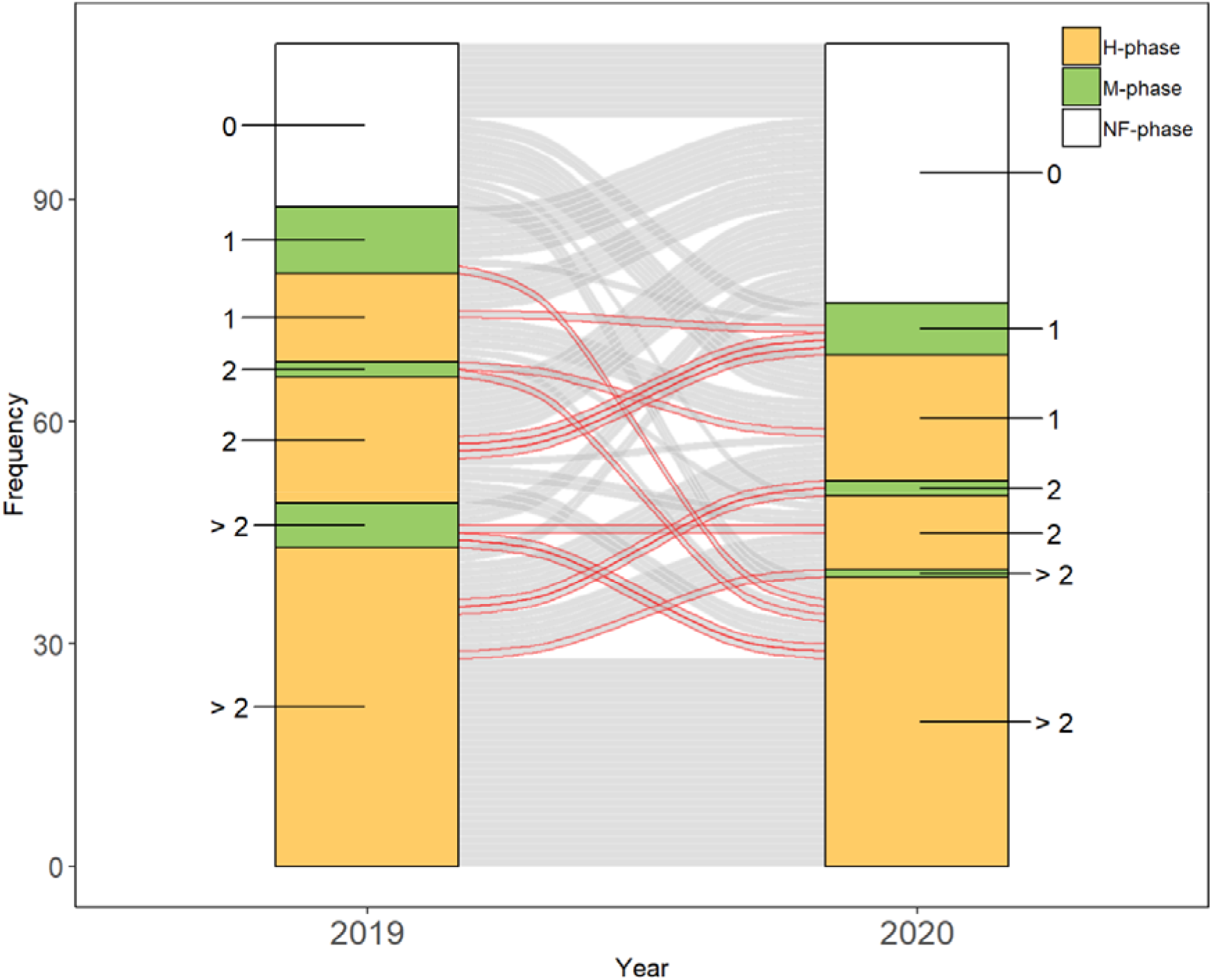
Flow diagram showing the changes of gender phase in marked individuals over two years in Population S1 (*N* = 111). The red lines represent changes in gender between an M-phase and an H-phase (*N*= 13, 12%). Individuals were separated into seven gender classes according to their gender phase (non-flowering, NF-phase; male, M-phase; hermaphrodite, H-phase) and the number of flowers (non-flowering, one-flowered, two-flowered, and more-than-two-flowered).

### Phenology and mate availability at the population level

In general, we found that larger individuals were more likely to flower than smaller individuals (Figure S3). We assessed flowering phenology and allocation in detail in Population LM in 2018. Flowering began on 29 May and ended on 29 June 2018 (149-180 on Julian day; Figure 2). Population LM comprised 625 individuals. In total, 899 flowers were produced during the season, of which 691 were phenotypically hermaphroditic and 208 were phenotypically male flowers. There were 111 individuals with only one male flower (18% of the population), 10 with multiple male flowers (1%), 342 with only one hermaphroditic flower (55%), and 162 with both male and hermaphroditic flowers (26%; Table 2). Hermaphroditic flowers (*N* = 103) from the subsampled individuals had a mean (± SD) of 184.8 ± 84.7 pistils and 202.3 ± 39.2 stamens, the latter being similar in number to those in male flowers (209.9 ± 38.3, *N* = 41; *t*-test: *P* = 0.3; Figure 2A and B). Such a pattern of pistil and stamen numbers is reflected in a bimodal distribution of the functional gender of the subsampled individuals (Figure 2C).

**Figure 2.**
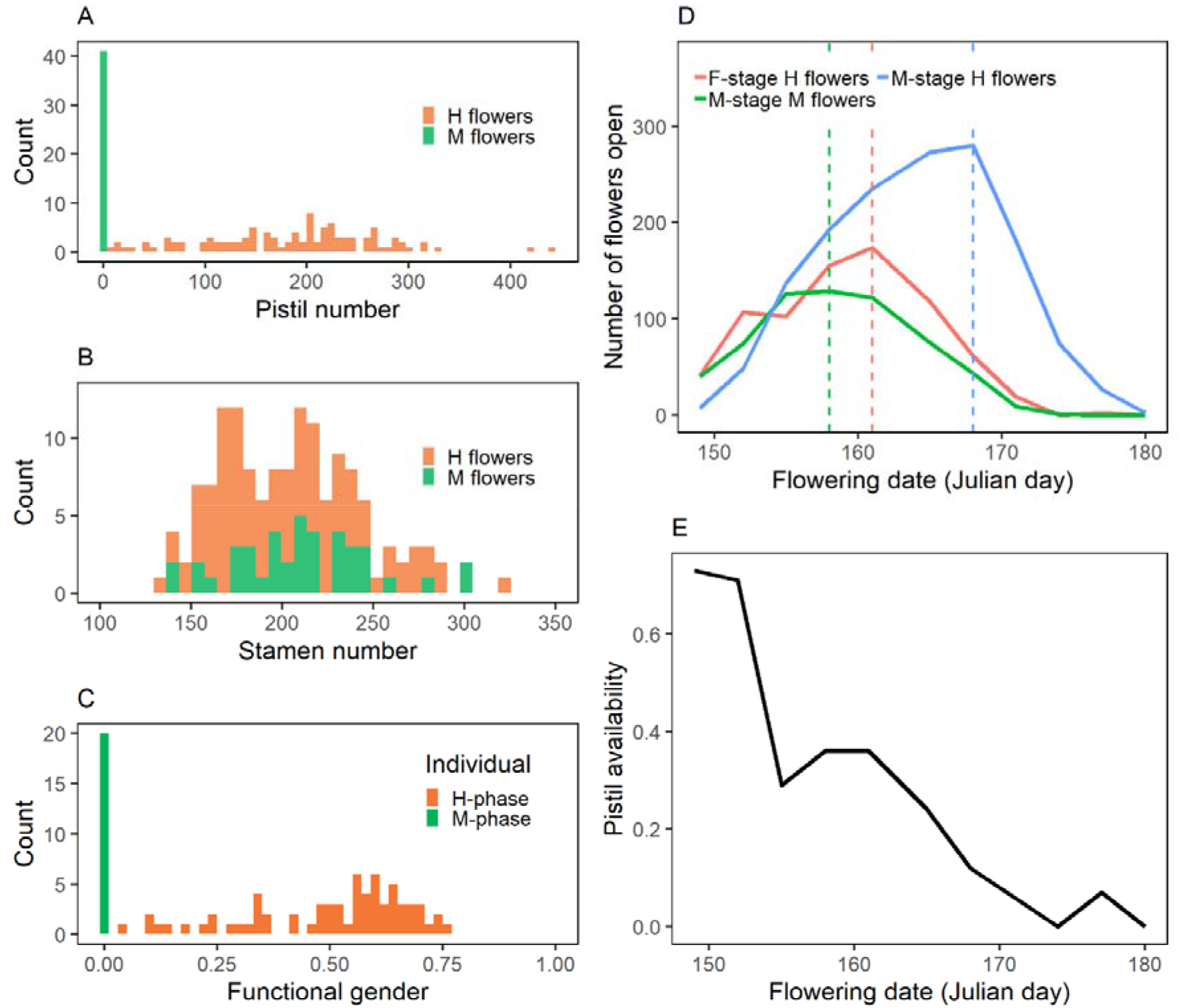
Histograms of the pistil number (A) stamen number (B) and functional gender (C) of sampled flowers, number of flowers at male and female stages (D), and pistil availability (E) across the flowering season in Population LM. ‘M flowers’ and ‘F flowers’ refer to phenotypically male and hermaphroditic flowers, respectively. ‘M-phase’ and ‘H-phase’ individuals refer to individuals flowering with only male or with hermaphrodite (and potentially male) flowers. ‘F-stage’ and ‘M-stage refer to flowers in their female or male stages (along their protogynous progression). (B) and (D) Phenotypically hermaphroditic (hermaphrodite flowers, *N* = 103) and male (Male flowers, *N* = 41) flowers are colored orange and green, respectively. The distributions of the pistil number (A) and functional gender (C) are bimodal whereas that of the stamen number (B) is unimodal. (D) Male flowers have only a male stage (M-stage M flowers, green line) while hermaphroditic flowers are first in their female stage (F-stage H flowers, orange line) and then in their male stage (M-stage H flowers, blue line). Note that the flowering peak for male flowers (green dashed line) is earlier than that for male-stage hermaphroditic flowers (blue dashed line). (E) Pistil availability, which reflects the ratio of available pistils to stamens at the population level, drops toward zero at the end of the flowering season.

Male and hermaphroditic flowers had the same longevity but differed in their phenology, with implications for the distribution of mate availability. Specifically, both male and hermaphroditic flowers lasted on average seven days, with hermaphroditic flowers spending three days in their female stage and four days in their male stage. The flowering peak for male flowers largely coincided with the peak for the female stage of hermaphroditic flowers. Male flowers thus preceded the peak for the male stage of hermaphroditic flowers (Figure 2D). Female mate availability dropped from 0.73 to zero over the course of flowering (Figure 2E), i.e., intra-sexual competition for siring success was low at first and increased over time. The observed decline in pistil availability over the flowering season in Population LM was also found in all four of the populations sampled in the subsequent year (2019; Figure S4).

### Size- and time-dependent SA, and functional gender

The number (proportion) of subsampled individuals with only one male flower, multiple male flowers, one hermaphroditic flower, and a mixture of male and hermaphroditic flowers were 17 (0.19), 3 (0.03), 39 (0.44), and 29 (0.33), respectively (Table 1). The absolute allocation to female and male functions among individuals varied from 0 to 0.37 g and from 0.009 to 0.09 g, respectively. Absolute allocation increased with size (Figure 3A; Table S4), but the slope of allocation on size was greater for the female function (95% CI of the coefficient: 0.032 - 0.065) than the male function (95% CI of the coefficient: 0.006 - 0.012; interaction between sex function and size: *P* < 0.01). This pattern was confirmed for the other four populations sampled in 2019 (Figure S3).

**Table 1.** Composition and number of flowers of individuals flowering as only male or as a hermaphrodite in Population LM. Shown are the number and proportion of individuals in each given phase, as well as the number and proportion that were subsampled for measurements. Note that the phenotypic gender of individuals producing only male flowers was classified as male (M-phase individuals) while that of individuals producing only hermaphrodite or a mixture of male and hermaphrodite flowers was classified as hermaphrodite (H-phase individuals).

|  | *N* | Proportion | Subsampled | Proportion |
| --- | --- | --- | --- | --- |
| 1-flowered M-phase | 111 | .18 | 17 | .19 |
| 1-flowered H-phase | 342 | .55 | 39 | .44 |
| 2-flowered M-phase | 8 | .01 | 3 | .03 |
| 2-flowered H-phase | 104 | .17 | 17 | .19 |
| >2-flowered M-phase | 2 | .003 | 0 | 0 |
| >2-flowered H-phase | 58 | .09 | 12 | .14 |
| Total | 625 | 1 | 88 | 1 |

**Figure 3.**
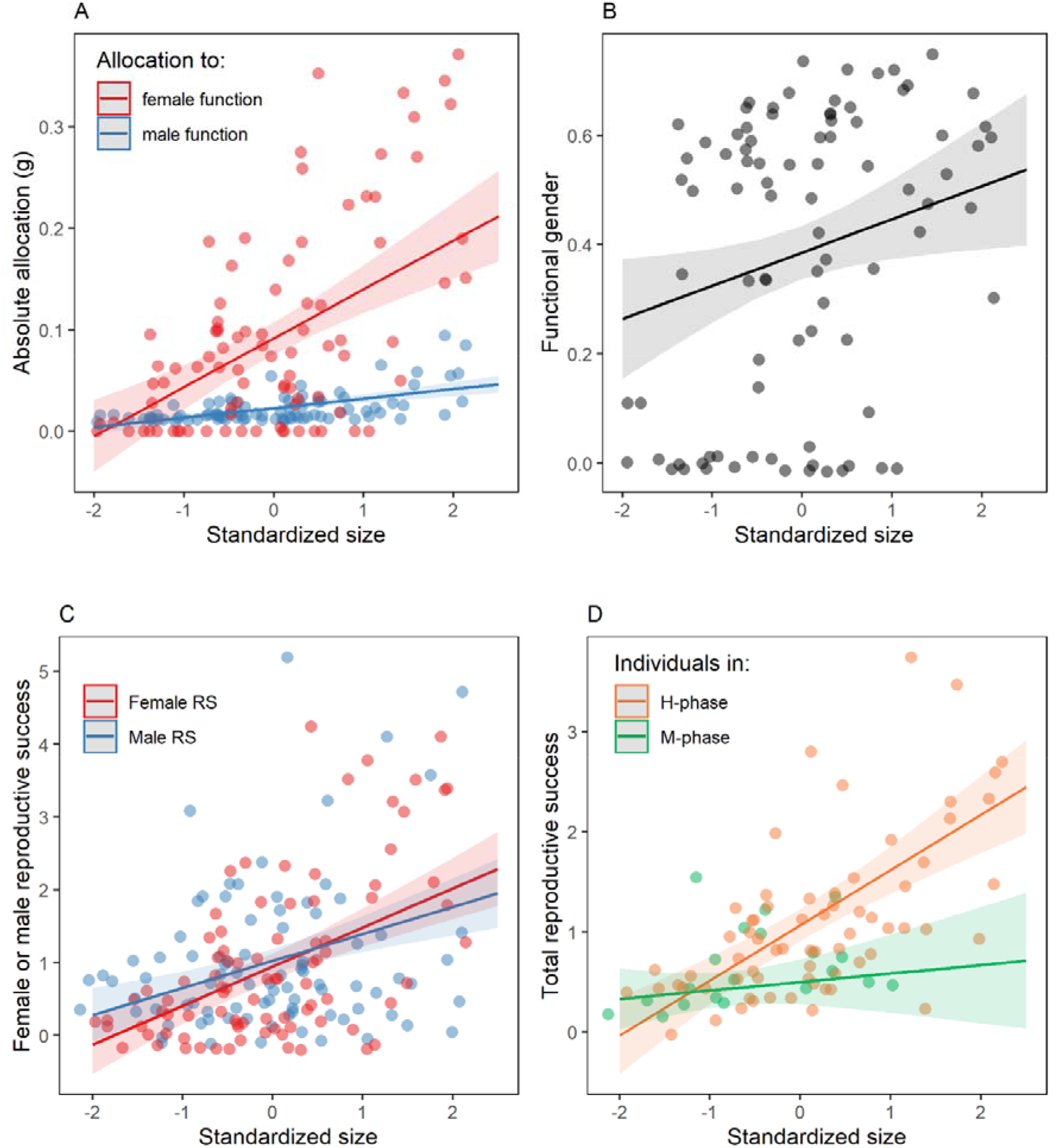
The effect of plant size on: (A) sex allocation to female and male functions (both slopes > 0, *P* < 0.001; female slope > male slope, size x sex interaction *P* < 0.001); (B) functional gender, calculated as femaleness (slope > 0, *P* < 0.01); (C) RS for individuals via their male and female functions (both slopes > 0, *P* < 0.001; female slope > male slope, size x sex interaction *P* < 0.05); and (D) total RS for male-phase and hermaphrodite-phase individuals (both slopes > 0, *P* < 0.001; H-phase slope > M-phase slope, size x gender interaction *P* < 0.001), as predicted by models fitted to data from Population LM. See text for details. ‘M-phase’ and ‘H-phase’ individuals refer to individuals flowering with only male or with hermaphrodite (and potentially male) flowers. Each individual contributes two points representing its female and male allocation in (A) and its relative female and male RS in (C). The 95 % confidence interval of the estimates is shown around the regression lines.

Absolute allocations also increased with flowering date (Figure 4A; Table S4), with the slope again larger for female function (95% CI of the coefficient: 0.019 - 0.051) than for male function (95% CI of the coefficient: -0.004 - 0.002; interaction between sex function and flowering date: *P* < 0.01). Functional gender among the sampled individuals ranged from 0 to 0.75 (where 0 represents pure male and 1.0 represents pure female; see also Figure 2C), increasing with size (*P* < 0.01; Figure 3B; Table S5) and flowering date (*P* < 0.001; Figure 4B; Table S5). Small and early-flowering individuals tended to be in a male-phase, i.e., with a functional gender of zero (Figure S5).

**Figure 4.**
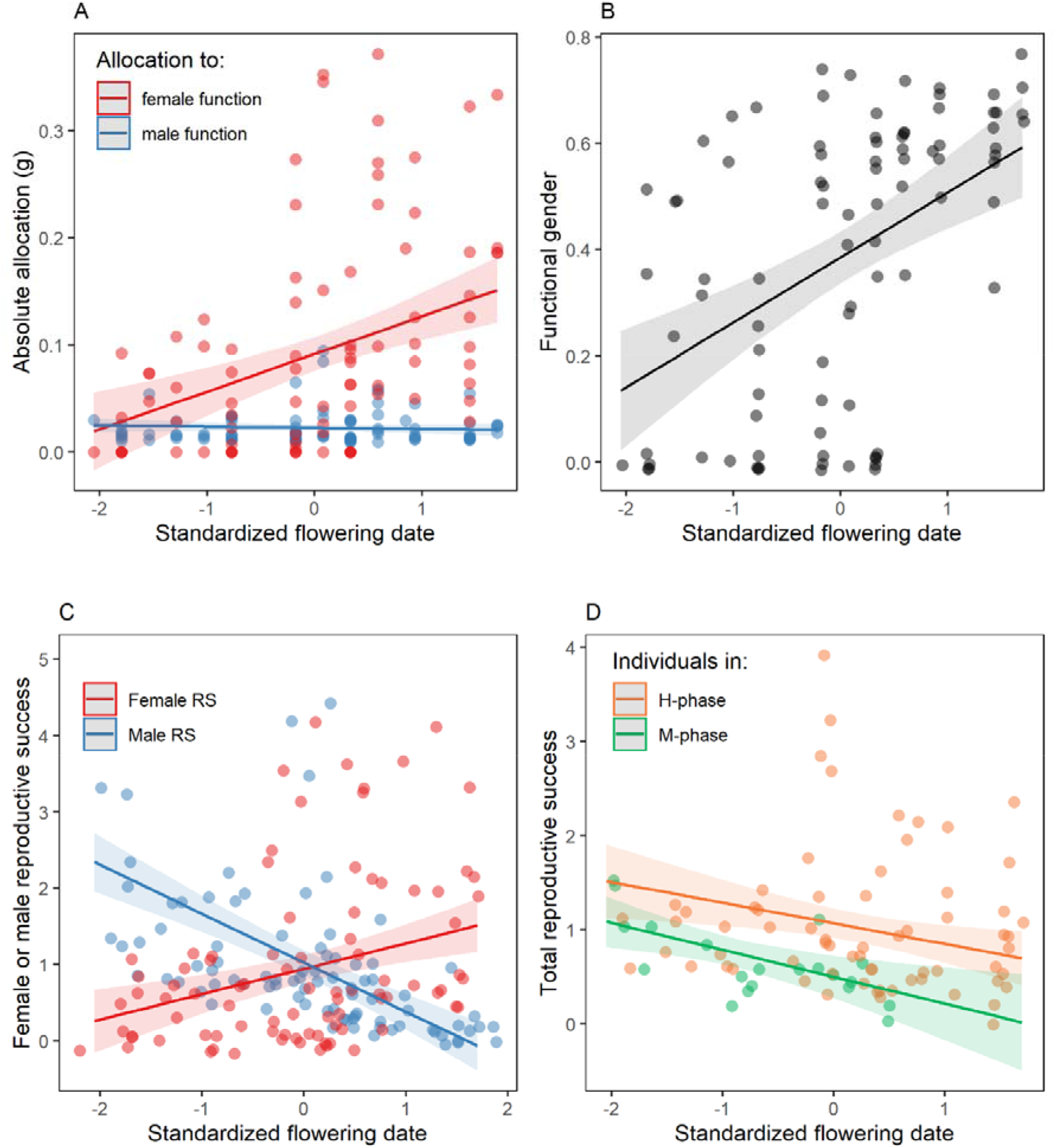
The effect of mid flowering date on: (A) sex allocation to female and male functions (female slope > 0, *P* < 0.001; flowering date x sex interaction *P* < 0.001); (B) functional gender, calculated as femaleness (slope > 0, *P* < 0.001); (C) RS for individuals via their male and female functions (female slope > 0, male slope < 0; flowering date x sex interaction *P* < 0.001); and (D) total RS for Male-phase and Hermaphrodite-phase individuals (both slopes < 0, *P* < 0.01; flowering date x gender interaction *P* > 0.05), as predicted by models fitted to data from Population LM. Each individual contributes two points representing its female and male allocation in (A) and its relative female and male RS in (C). ‘M-phase’ and ‘H-phase’ individuals refer to individuals flowering with only male or with hermaphrodite (and potentially male) flowers. The 95 % confidence interval of the estimates is shown around the regression lines.

### Size- and time-dependent prospective seasonal RS

Relative RS increased with plant size (Figure 3C; Appendix table S4; *P* < 0.001), but the slope was slightly steeper for female RS (95% CI of the coefficient: 0.35 to 0.72) than for male RS (95% CI of the coefficient: 0.2 to 0.54; interaction between size and sex function: *P* < 0.05). The relationship between flowering date and RS differed between male and female functions (interaction: *P* < 0.001). Female RS increased with flowering date (Figure 4C; Appendix table S4; 95% CI of the coefficient: 0.16 to 0.51), whereas male RS decreased with flowering date (95% CI of the coefficient: -0.81 to -0.48). There was a three-way interaction among size, flowering date, and sex function (Table S4; Figure S6; *P* < 0.05).

Total RS increased with size (Figure 3D; Table S6; *P* < 0.001), but the slope was steeper for hermaphrodite-phase (95% CI of the coefficient: 0.38 to 0.73) than for male-phase individuals (95% CI of the coefficient: -0.12 to 0.29; interaction between size and gender: *P* < 0.001). Total seasonal RS decreased with flowering date for both hermaphrodite-phase and male-phase individuals (Figure 4D; Table S6; *P* < 0.01; interaction between flowering date and gender: *P* > 0.05).

### Phenology of andromonoecious individuals

For individuals expressing andromonoecy, i.e., individuals producing both male and hermaphroditic flowers in the same season, the first flower had a highest probability of being a male flower, at a time of maximum female mate availability (Figure 5).

**Figure 5.**
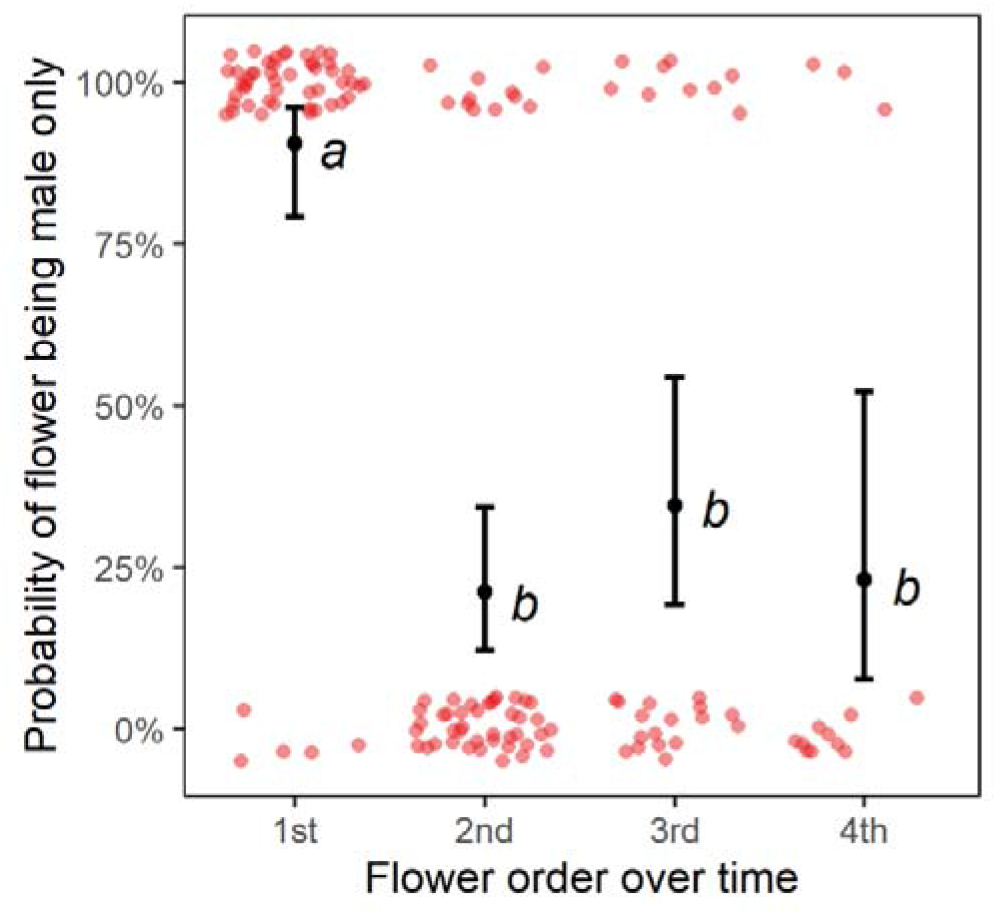
Probability that individual flowers are male only (have stamens but not pistils) as a function of their opening order on an individual. The first flowers to open on multi-flowered individuals have a much higher probability of being male than flowers opening subsequently. Data are from Population LM and are based on 144 flowers over 54 individuals. The probabilities are shown with the 95 % confidence interval predicted by the model (see text for details). Letters indicate means that are not significantly different from one another, based on a Tukey test (*P* > 0.05).

## Discussion

We assessed the effect of plant size and the within-season timing of flowering on SA and prospective RS in the andromonoecious plant *P. alpina*. Our results corroborate theories on size-dependent SA and illustrate the likely influence of resource status on gross flowering decisions made by perennial plants over the course of their lives. In particular, they help to explain both the expression of gender diphasy in *P. alpina* as well as the evolution of andromonoecy in species with a strong temporal separation of the male and female functions in their flowers.

### *The mating system and inbreeding depression in* P. alpina

Results from our hand-pollination treatments indicate that *P. alpina* is self-compatible and that there are no differences in the ability of pollen from male and hermaphrodite flowers of *P. alpina* to sire progeny upon pollination. Although self-incompatibility systems are common in Ranunculaceae (Allen and Hiscock 2008), most of the species in the genus *Pulsatilla*, including *P. alpina* in this study, are self-compatible (Jonsson et al. 1991; Lindell 1998). No difference in the siring ability of pollen from male and hermaphroditic flowers further justifies our estimate of male RS, which conforms to the general pattern found in other andromonoecious species (Solomon 1985; Huang 2003; Cuevas and Polito 2004; Dai and Galloway 2012).

The finding that artificially self-pollinated flowers of *P. alpina* produced fewer seeds than those pollinated with outcross pollen points to the expression of a degree of early-acting inbreeding depression in the species. Our estimate of inbreeding depression at the seed stage is 0.15, which should be interpreted as the low bound given that the inbreeding depression may expression at the later stages and that the inbreeding depression is in general strong in perennial plants (Angeloni et al. 2011). Nonetheless, inbreeding depression will most likely reduce the male RS estimated by our mass-action model of the individuals with multiple flowers via geitonogamy (though not for individuals with single flowers, which we assume are fully outcrossed).

### Size- and resource-dependent SA

*P. alpina* is a perennial herb that stores its resources over winter and protects its meristems for future growth in an underground rhizome. We did not measure the size of the rhizome of the plants sampled in this study, but we suppose that above-ground plant size strongly reflects rhizome size as a function of both age and factors that impact resource gains (through photosynthesis) and resource losses (e.g., through flowering, fruiting and/or as a result of herbivory) in the previous growing season(s). The observation that small plants were less likely to flower than large plants is consistent with this supposition, as is the fact that small plants were less likely to flower in two consecutive years than were larger plants.

Our results are also consistent with the notion that flowering through the female function (in *P. alpina*, this means adopting a hermaphroditic rather than a male phase in a given season) places a heavier burden on a plant’s resources than flowering just through its male function (Schlessman 1991; Zhang et al. 2014; Bialic-Murphy et al. 2020): smaller plants were more likely to produce a single male flower, whereas larger plants produced more flowers and flowers with both male and female functions. Indeed, we found that larger individuals of *P. alpina* made a greater absolute and relative allocation to their female function than smaller plants, with a significantly steeper positive slope of absolute allocation to the female than male function. Accordingly, larger plants also expressed a more female functional gender than small plants. A similar size-dependent SA strategy has been found in other insect-pollinated perennials (reviewed in de Jong and Klinkhamer 2006) and conforms to predictions of theory on size-dependent SA in simultaneously hermaphroditic plants (Klinkhamer et al. 1997; Cadet et al. 2004).

### Gender diphasy with small functional males

An important feature of the reproductive strategy of *P. alpina* is that its floral SA shows both variation on a continuum, particularly with different numbers of pistils produced in its flowers, but also discrete variation involving the production of either bisexual hermaphroditic flowers or male flowers. Significantly, male flowers do not represent just the extreme end of a male-female continuum, but rather one of two quite different modes of floral allocation, with flowers producing no pistils at all or a few hundred pistils. This means that small individuals that only produce one or (rarely) two male flowers have adopted a fully male strategy for the season, whereas larger individuals flower as hermaphrodites. To the extent that size reflects age, this dichotomy represents a strategy of gender diphasy, where individuals transition from one allocation mode to another (Schlessman 1988). Indeed, our transition matrix reveals that about a tenth of all individuals changed their gender between consecutive seasons (see also Table S2).

Although gender diphasy in perennial plants most commonly involves a shift in gender expression from male to female, many species share the strategy displayed by *P. alpina* of shifting between male- and hermaphrodite-phases (Freeman et al. 1980; Schlessman 1988), e.g., *Lilium apertum* (Zhang et al. 2014), *Lloydia oxycarpa* (Niu et al. 2017), *Panax trifolium* (Schlessman 1991), and *Tulipa pumila* (Astuti et al. 2020). In these species, male-phase individuals are usually small and produce only one or few flowers, and it is generally thought that the small individuals in the male-phase likely contribute little to their lifetime fitness (Charlesworth 1984; Zhang and Jiang 2002). Indeed, studies on gender diphasic species have typically failed to show any advantages in male RS of male-phase over hermaphrodite-phase individuals in terms of pollen production, flower size, pollinator visitation rate, or pollen siring ability (Peruzzi et al. 2012; Zhang et al. 2014; Niu et al. 2017; Astuti et al. 2020). However, in most of these cases, the male flowers are smaller and produce less pollen compared to hermaphroditic flowers (Zhang et al. 2014; Niu et al. 2017; Astuti et al. 2020). This contrasts with the male flowers of *P. alpina*, which tend to produce the same number of stamens as hermaphroditic flowers. We discuss the significance of this observation in the next section.

A strategy of size-dependent SA needs to be understood in terms of both the costs and the benefits of flowering through a particular sexual function. In their model of size-dependent SA, Zhang and Jiang (2002) suggested that small individuals allocating only to male function could maximize their lifetime fitness by keeping their reproductive effort low and thereby enhancing their survival and RS in the following seasons. They argued that this should be especially so when the marginal cost of the female function is substantially higher than the male function. To some extent, this scenario would seem to apply to *P. alpina*. In a previous study (Chen and Pannell 2022), they found that individuals of *P. alpina* bear a particularly heavy cost of their female function in terms of the elongated floral stalks produced for hermaphroditic flowers but not male flowers – presumably as an adaptation for seed dispersal by wind. Small and resource-limited individuals of *P. alpina* might avoid allocation to female function and the need to produce costly floral stalks by producing male flowers partly as a strategy to enhance inter-seasonal survivorship. However, our results suggest that desisting from female allocation in *P. alpina* also has an alternative or additional explanation in terms of capitalizing on opportunities for siring success.

### Implications of the timing of SA for female versus male RS

Our study stands out by having estimated not only the allocation to male and female functions of individuals of different sizes and resource status but also the time-dependent opportunities for mating and RS over the course of the flowering season. Because flowers of *P. alpina* are strongly protogynous, opportunities for mating, estimated in terms of the number of receptive pistils available per stamen, declined steeply during the course of the flowering season (see Austen et al. 2015 for a review of how the other factors may affect this relation). We estimated female RS for each plant in terms of the number of seeds it produced during the season. Prospective male RS of individuals was determined by integrating their prospective siring success in terms of a mass action model of mating over the period of their flowering, accounting for both their stamen production and the numbers of pistils for which they might compete to pollinate within successive time windows. We could thus estimate the prospective seasonal RS for each individual both as a function of its size and also in terms of when during the season it was likely to have been most successful through its male function.

Our results indicate that larger plants had greater prospective seasonal RS through both their male and female functions, but the relationship with size was significantly steeper for the female function, with an intersection between the two sex functions at mid-size. This indicates that male RS contributes relatively more to the total seasonal RS of small plants whereas, after crossing a certain size threshold, the contribution of female RS likely exceeds that of the male function – a pattern that is reflected in the overall greater female gender of larger plants. The reproductive allocation strategy of *P. alpina* thus conforms to the size-advantage hypothesis (Ghiselin 1969; Zhang and Jiang 2002), not only in terms of its gender diphasy, discussed above, but also in terms of the quantitative variation in gender among hermaphrodites of different sizes but flowering together in the same season.

An important implication of our results is that the dynamic nature of female mate availability over the course of the flowering season meant that plants flowering early tended to have a male-biased gender, while those flowering later had a female-biased gender. Moreover, because of the high availability of pistils to be pollinated in the early season, small plants that produce only male flowers (see above) had high prospective siring success despite their small size, i.e., their male-only allocation is probably not just an outcome of resource constraints facing small individuals, as often supposed for other species (Zhang and Jiang 2002; Peruzzi et al. 2012), but also an adaptation to capitalize on mating opportunities. Indeed, our results indicate that although small male-phase individuals obviously gained no fitness via their female function, the prospective total seasonal RS of some male-phase individuals is likely to have been higher than that of some hermaphrodite-phase individuals of similar size.

Most studies have not considered the effect of both plant size and flowering phenology (and patterns of mate availability) jointly (but see Schlessman et al. 1996; Schlessman and Graceffa 2015). Consequently, the potentially high contribution to lifetime fitness of plants via their male function when small, and thus the functional significance of small males in gender diphasic species (Kudo and Maeda 1998; Zhang and Jiang 2002), may hitherto have been under-appreciated. According to our reasoning, we should expect male-phase individuals in diphasic populations to be early flowering in protogynous species, as in *P. alpina*, and late flowering in protandrous species. Indeed, in protandrous dwarf ginseng (*Panax trifolium*), male-phase individuals were found to flower later in the season compared to hermaphrodite-phase individuals and were highly synchronized with the phenology of the female-stage of hermaphroditic individuals (Schlessman et al. 1996). These patterns are generally consistent with ideas from SA theory and the dynamic nature of the mating environment in dichogamous species (Brunet and Charlesworth 1995; Austen and Weis 2014) and are coherent with other empirical studies on protogynous species (Huang et al. 2004; Guitián 2006).

### Andromonoecy as the resolution of intersexual conflict due to dichogamy

Our analysis of SA and components of prospective RS for individuals of *P. alpina* in the context of their flowering phenology also points to an explanation for the evolution of andromonoecy in dichogamous species. Andromonoecy is a heteromorphism involving the production of male and hermaphrodite flowers by the same individual. Although small individuals of *P. alpina* produce only male flowers, over the course of their lives all individuals likely produce both male and hermaphroditic flowers, so the sexual system can be considered andromonoecious. Our results suggest that the male flowers of large individuals likely promote male RS in the same way as the early flowering of small male-phase individuals, notably by resolving intersexual conflict faced by individuals with bisexual flowers.

As our analysis of phenology clearly shows, plants stand to achieve substantial siring success by dispersing pollen early in the flowering season. But because *P. alpina* is strongly protogynous, individuals producing bisexual flowers must delay the onset of their male function until they have passed through the female stage. This immediately suggests that andromonoecy in *P. alpina* might be interpreted as a strategy to advance the timing of the male function of individuals by suppressing their female function in some flowers. The resolution of the sexual conflict within flowers through the production of all-male flowers rather than via a modification of the dichogamy responsible for the conflict is consistent with the fact that species in the genus *Pulsatilla* are all protogynous and that patterns of dichogamy tend to be much more phylogenetically conserved than phenology and could thus be viewed as a phylogenetic constraint (Lloyd and Webb 1986; Jonsson et al. 1991; Routley et al. 2004).

The function of male flowers in andromonoecious species has attracted substantial speculation (Tomaszewski et al. 2018), but empirical evidence for the various ideas put forward remains ambiguous. For example, male flowers in *Passiflora incarnata* sired on average twice as many seeds as hermaphroditic flowers, largely as a result of greater pollen production and less self-pollen deposition (Dai and Galloway 2012), and male RS in *Solanum carolinense* increased with the proportion of male flowers but not with the total number of flowers, likely because the absence of pistils in male flowers allowed bumblebees to remove pollen more efficiently (Elle and Meagher 2000). In contrast, Podolsky (1993) found that hermaphroditic flowers in *Besleria triflora* dispersed substantially more pollen than male flowers over an average flower’s lifetime, implying that male flowers contributed relatively little directly to male RS. Similarly, male flowers in *Anticlea occidentalis* promoted female mating quality in terms of outcrossing rate and mate diversity but did not affect male RS in terms of seed sired (Tomaszewski et al. 2018).

Our study illustrates the insights that can be gained by studying the life history and phenology of plants jointly rather than independently. Moreover, by calling attention to the potentially high contribution to seasonal male RS made by male flowers produced when mate availability is particularly high, and by drawing a link with gender diphasy, our study exposes a simple but largely overlooked explanation for the evolution of andromonoecy (Pellmyr 1987; Schlessman 2010). Indeed, both gender diphasy and andromonoecy have been found in many dichogamous species in Ranunculaceae (Pellmyr 1987; Lindh 2017), Liliales (Peruzzi 2012), and Apiales (Schlessman 2010), and Schlessmann (2010) has argued that the flowering order of male and hermaphroditic flowers in such species should depend on whether they are protandrous or protogynous. In common with *P. alpina*, these species are also often perennial herbs with underground storage organs, a relatively short flowering season, and a relatively high cost of female function associated with either the dispersal of seeds by wind from costly inflorescence stalks or by animals attracted to costly fleshy fruits.

Taken together, our results show that resource status affects the absolute and relative resource allocation to female function in *P. alpina*, while the timing of flowering likely determines RS through its male function. This is consistent with the general view that female RS is commonly limited by resources, whereas male RS is limited by mate availability (Bateman 1948; Charnov 1979). The gender-diphasic and andromonoecious strategy displayed by *P. alpina* means that individuals can adjust their SA in response to resource status and mate availability at both the individual level, from male-phase to hermaphrodite-phase individuals, and the flower level, from phenotypically male flowers to hermaphroditic flowers – a dichotomy not predicted by theory for SA that consider size and the timing of flowering independently (Brunet and Charlesworth 1995; Klinkhamer et al. 1997). As a consequence, individuals may avoid investing in their female function when resources are limited, consistent with predictions for the evolution of andromonoecy (Spalik 1991; de Jong et al. 2008), and by producing male flowers early in the season when mate availability is high.

## Supplementary Materials

**Table S1.** Descriptions of the study populations and the studies conducted in them from 2018 to 2020.

| Population | Location | GPS position | Altitude<br>(a.s.l.) | *N* | Flower<br>season<sup>+</sup> | Habitat | Note | SA and RS<br>estimates<br>(2018) | Reproductive<br>biology<br>(2018) | Generality of SA<br>(2019) | Gender change<br>(2019-2020) |
| --- | --- | --- | --- | --- | --- | --- | --- | --- | --- | --- | --- |
| LM | Les Mosses | 46°23'57"N<br>7°04'52"E | 1694 | About 600 | E | enclosed<br>grassland | plant size<br>mostly small | O |  |  |  |
| LL1 | Lac Lioson | 46°23'08"N<br>7°07'23"E | 1951 | > 1,000 | E | open grassland | steep slope |  | O | O |  |
| LL4 | Lac Lioson | 46°22'57"N<br>7°07'11"E | 1983 | > 1,000 | L | open grassland |  |  |  | O |  |
| S1 | Solalex | 46°17'36"N<br>7°09'11"E | 1723 | > 1,000 | E | open grassland | population<br>fenced |  |  | O | O |
| S2 | Solalex | 46°16'37"N<br>7°09'32"E | 2122 | > 1,000 | L | open grassland | steep slope |  |  | O |  |

+ Flowering season indicates whether the population starts flowering in late May to early June (E) or late June to July (L).

**Table S2.** Description of phenotypic classes at different levels used in this study.

| Levels | Phenotypic classes | Description |
| --- | --- | --- |
| Individual-level | Male-phase individual | Individuals with male function only, i.e., only male flowers. |
|  | Hermaphrodite-phase individual | Individuals with both male and female functions, i.e., with hermaphroditic and potentially male flowers. |
| Flower-level | Male flower | Flowers with stamens only |
|  | Hermaphroditic flower | Flowers with both stamens and pistils |
| Intra-flower level | Female-stage flower | Hermaphrodite flowers with receptive pistils (the Female stage precedes the Male stage in hermaphrodite flowers). |
|  | Male-stage flower | Male or hermaphrodite flowers with dehiscent stamens. |

**Table S3.**
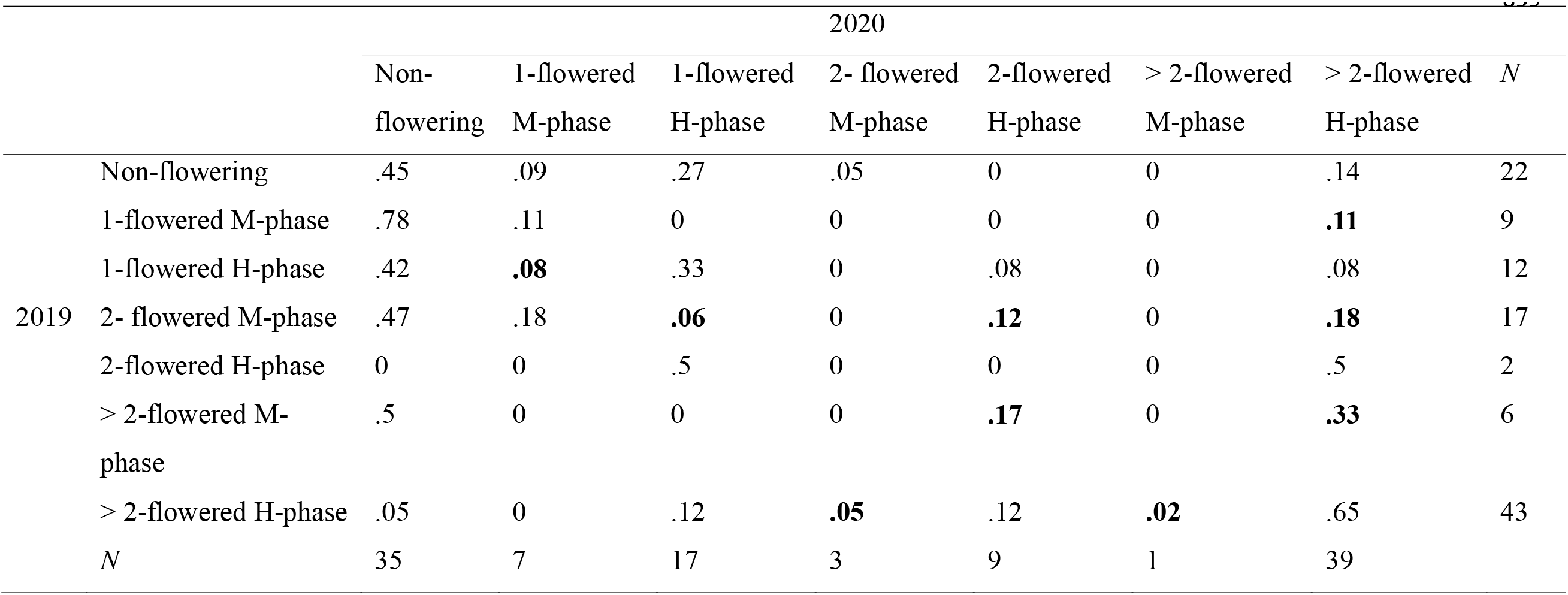
Transition matrix of marked individuals at different flowering states or phases between two years in population S1 (N = 111). The rates in bold indicate a transition between male-phase (M-phase) and hermaphrodite-phase (H-phase) individuals.

|  |  | 2020 |  |  |  |  |  |  | 899 |
| --- | --- | --- | --- | --- | --- | --- | --- | --- | --- |
| | | Non-flowering | 1-flowered<br>M-phase | 1-flowered<br>H-phase | 2- flowered<br>M-phase | 2-flowered<br>H-phase | > 2-flowered<br>M-phase | > 2-flowered<br>H-phase | $N$ |
| 2019 | Non-flowering | .45 | .09 | .27 | .05 | 0 | 0 | .14 | 22 |
|  | 1-flowered M-phase | .78 | .11 | 0 | 0 | 0 | 0 | **.11** | 9 |
|  | 1-flowered H-phase | .42 | **.08** | .33 | 0 | .08 | 0 | .08 | 12 |
|  | 2- flowered M-phase | .47 | .18 | **.06** | 0 | **.12** | 0 | **.18** | 17 |
|  | 2-flowered H-phase | 0 | 0 | .5 | 0 | 0 | 0 | .5 | 2 |
|  | > 2-flowered M-phase | .5 | 0 | 0 | 0 | **.17** | 0 | **.33** | 6 |
|  | > 2-flowered H-phase | .05 | 0 | .12 | **.05** | .12 | **.02** | .65 | 43 |
| $N$ | | 35 | 7 | 17 | 3 | 9 | 1 | 39 | |

**Table S4.** Summary table of the effects of plant size, flowering date, and sex function (male or female) on absolute sex allocation (SA) and reproductive success (RS). Absolute allocation was calculated as the dry mass of pistils (female function) and stamens (male function) of each individual. RS was estimated by the number of mature seeds and by a mass-function model for female and male functions respectively (see main text for details).

|  | Absolute<br>allocation (*gls*)<sup>+</sup> |  |  | Relative RS (*gls*)<sup>+</sup> |  |  |
| --- | --- | --- | --- | --- | --- | --- |
|  | df | LRT | *P* | df | LRT | *P* |
| Size | 1 | 4.77 | < .001 | 1 | 40.8 | < .001 |
| Flowering date | 1 | 16 | < .05 | 1 | 6.42 | < .05 |
| Sex function | 1 | 43.89 | < .001 | 1 | 0 | n.s. |
| Sex function x Size | 1 | 29.6 | < .001 | 1 | 4.44 | < .05 |
| Sex function x Flowering date | 1 | 20.02 | < .001 | 1 | 46.32 | < .001 |
| Size x Flowering date | 1 | .59 | n.s. | 1 | .62 | n.s. |
| Sex function x Size x Flowering<br>date | 1 | 4.6 | < .05 | 1 | 6.58 | < .05 |

Note—Individual identity was set in a compound symmetry structure in the two models to take into account the correlation of male and female sex function from the same individual.
+ Variance was allowed to vary between sex functions in the *gls* models (see main text for details).

**Table S5.**
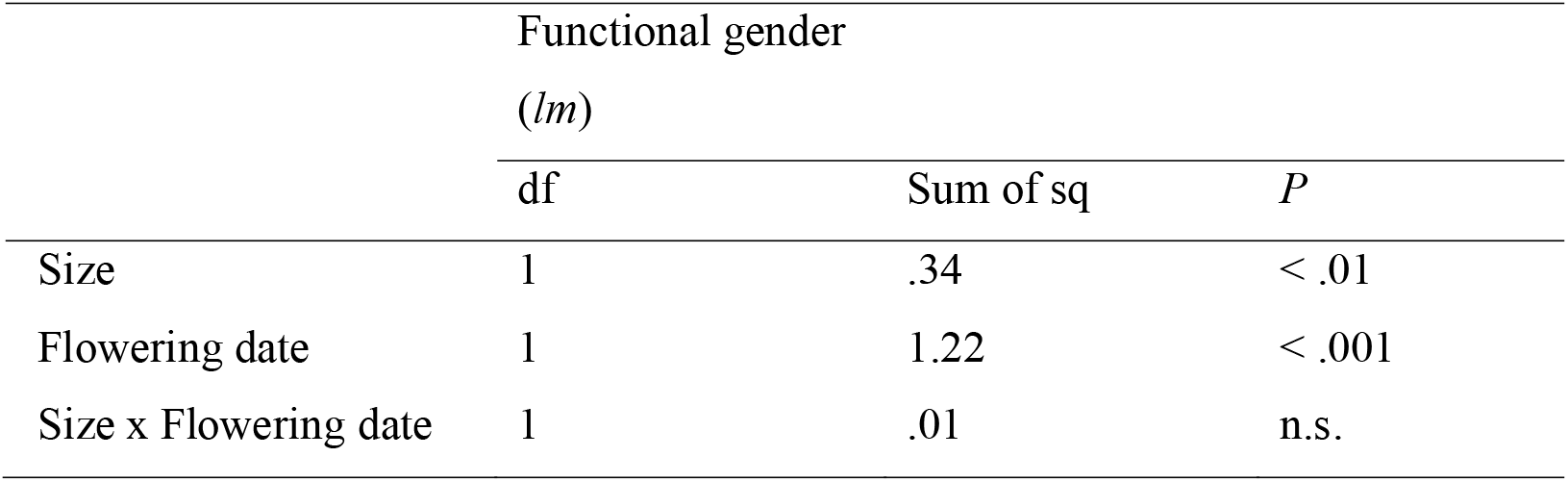
Summary table of the effects of size and flowering date on functional gender. Functional gender was calculated in terms of femaleness (see main text for details).

**Table S6.** Summary table of the effects of size, flowering date, and phenotypic gender (male phase or hermaphrodite phase) of individuals on relative total reproductive success. The total RS of the individuals was calculated as the sum of female and male RS (see main text for details).

|  | Relative total<br>reproductive<br>success (*gls*)<sup>+</sup> |  |  |
| --- | --- | --- | --- |
|  | df | LRT | *P* |
| Size | 1 | 24.79 | < .001 |
| Flowering date | 1 | 8.96 | < .01 |
| Phenotypic gender | 1 | 6.28 | < .05 |
| Phenotypic gender x Size | 1 | 13.07 | < .001 |
| Phenotypic gender x Flowering date | 1 | .756 | n.s. |
| Size x Flowering date | 1 | .02 | n.s. |
| Phenotypic gender x Size x<br>Flowering date | 1 | .57 | n.s. |

Note—+ Variance was allowed to vary between phenotypic genders in the *gls* model (see main text for details).

**Figure S1.**
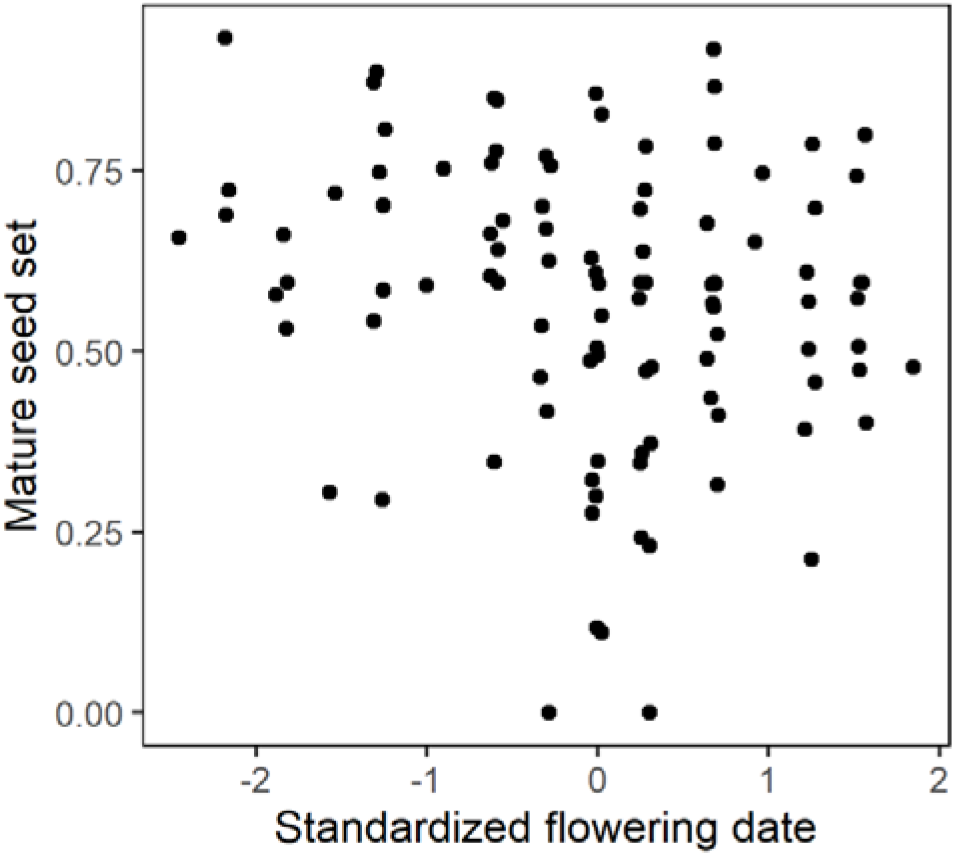
Mature seed set per flower as a function of the standardized flowering date in Population LM in 2018. The mature seed set of each flower is independent of the flowering date (*P* > 0.05). Each point represents one hermaphroditic flower (*N* = 103 flowers from 68 individuals). The results indicate that there is likely no change in the degree of pollen- and resource-limitation across the flowering season regarding reproduction via the female function.

**Figure S2.**
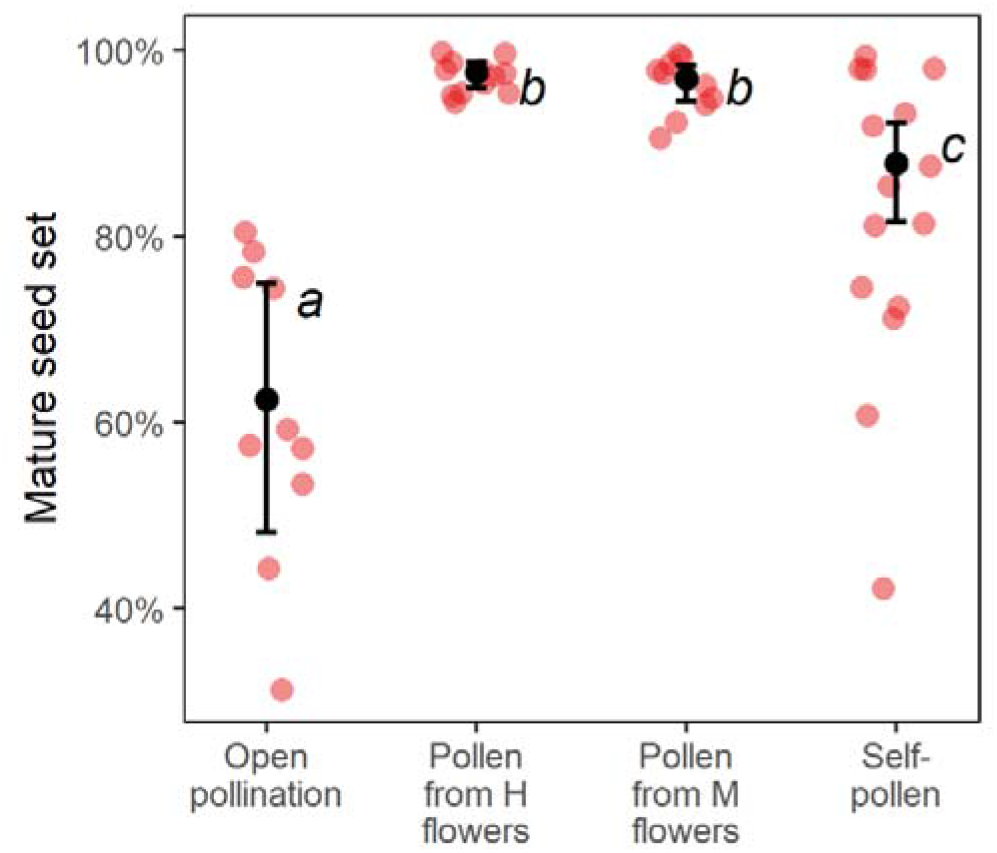
The effect of artificial pollination treatment on mature seed set in Population LL1. Pollination treatments included open pollination (*N* = 10), artificial outcrossing with pollen from hermaphrodite flowers (*N* = 12), outcrossing with pollen from male flowers (*N*= 10), and artificial selfing (*N* = 15). The mature seed sets are shown with the 95 % confidence interval predicted by a generalized linear mixed model. Letters indicate means that are not significantly different from one another, based on a Tukey test (*P* > 0.05).

**Figure S3.**
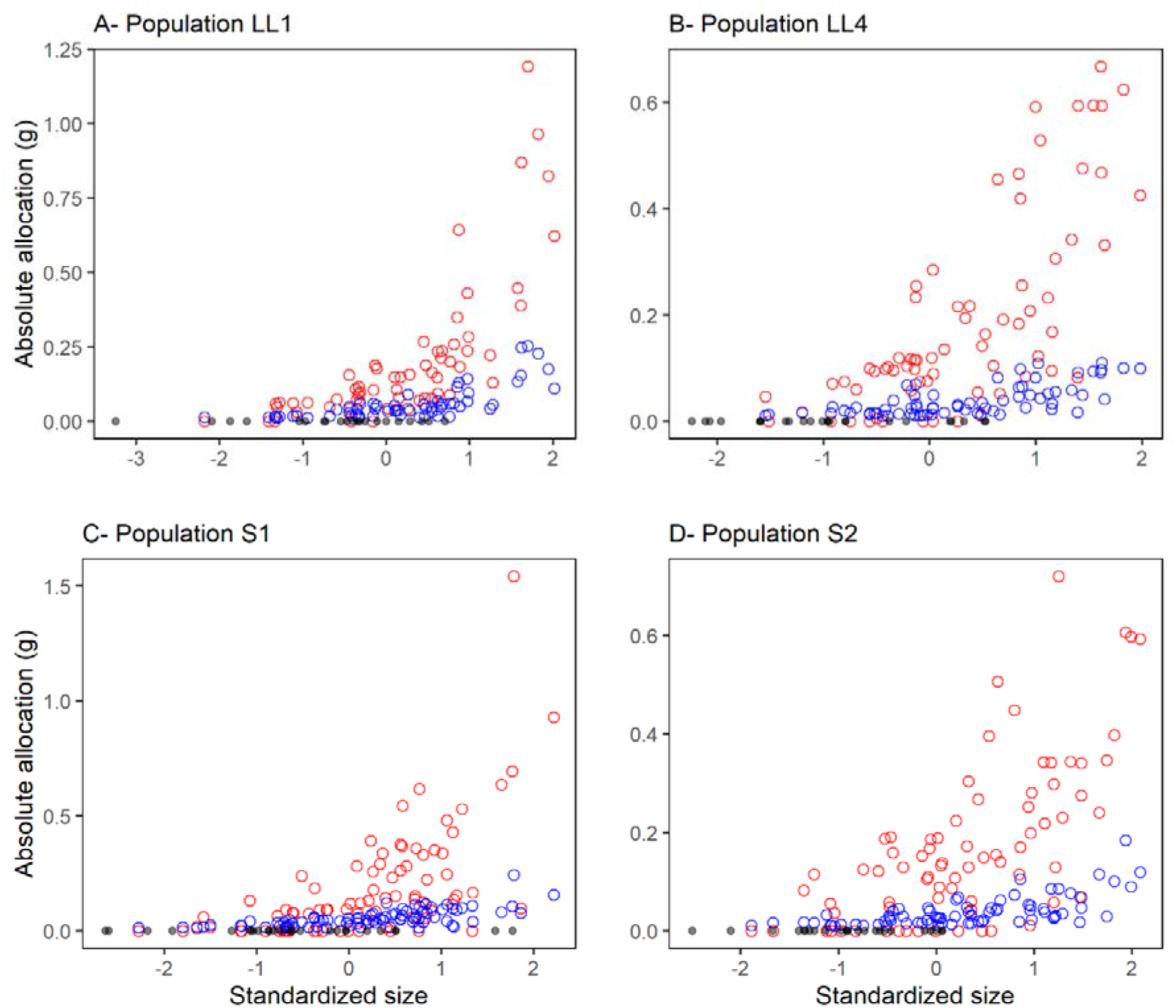
Plant size and absolute allocation to male and female functions in Populations LL1 (A), LL4 (B), S1 (C), and S2 (D). Each flowering plant is represented by two circles for its female (red circle) and male functions (blue circle). Non-flowering individuals are represented by grey points. The sample size of flowering and non-flowering individuals in Populations LL1, LL4, S1, and S2 are 61, 70, 82, and 75 and 21, 24, 26, and 24, respectively.

**Figure S4.**
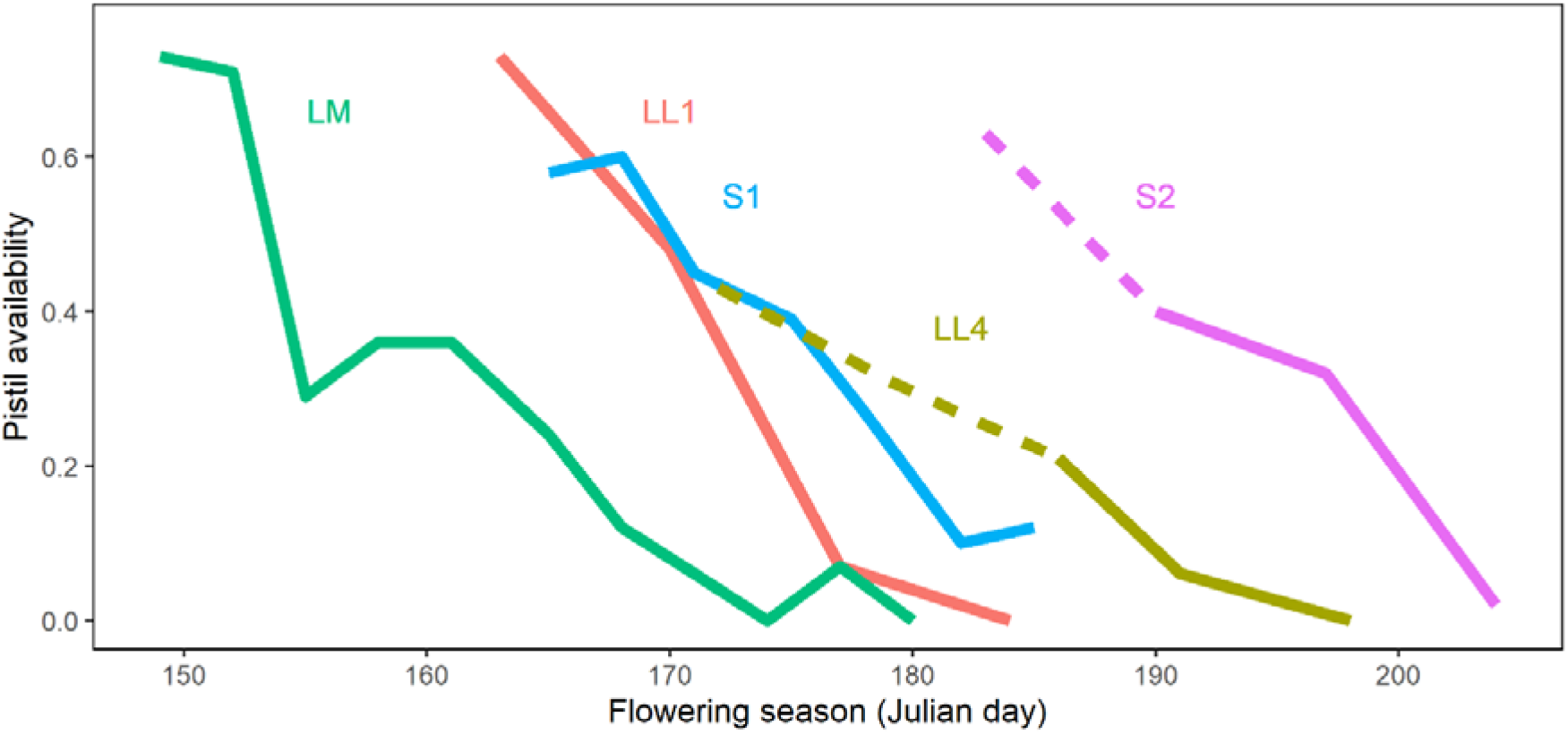
Changes in pistil availability across the flowering season in five populations. Pistil availability was monitored in the spring of 2018 (LM) and 2019 (LL1, LL4, S1, and S2). The observations in Populations LL4 and S2 were conducted from the peak of the flowering season rather than from the early flowering season; the dashed lines are thus extrapolations to the inferred starting of the flowering date based on the assumption of a linear decrease in pistil availability over the flowering season from the field data.

**Figure S5.**
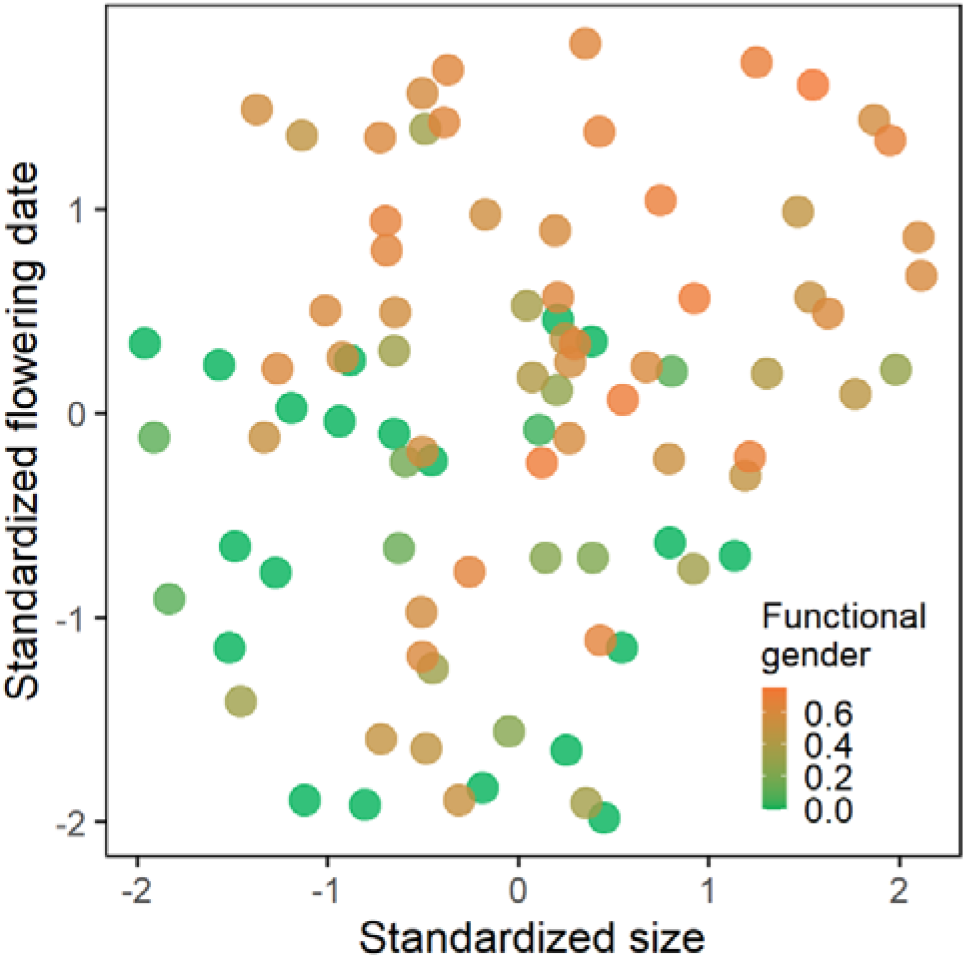
Effects of plant size and flowering date on functional gender of sampled individuals in Population LM. Each point represents one individual and the color of the point indicates its functional gender calculated in femaleness.

**Figure S6.**
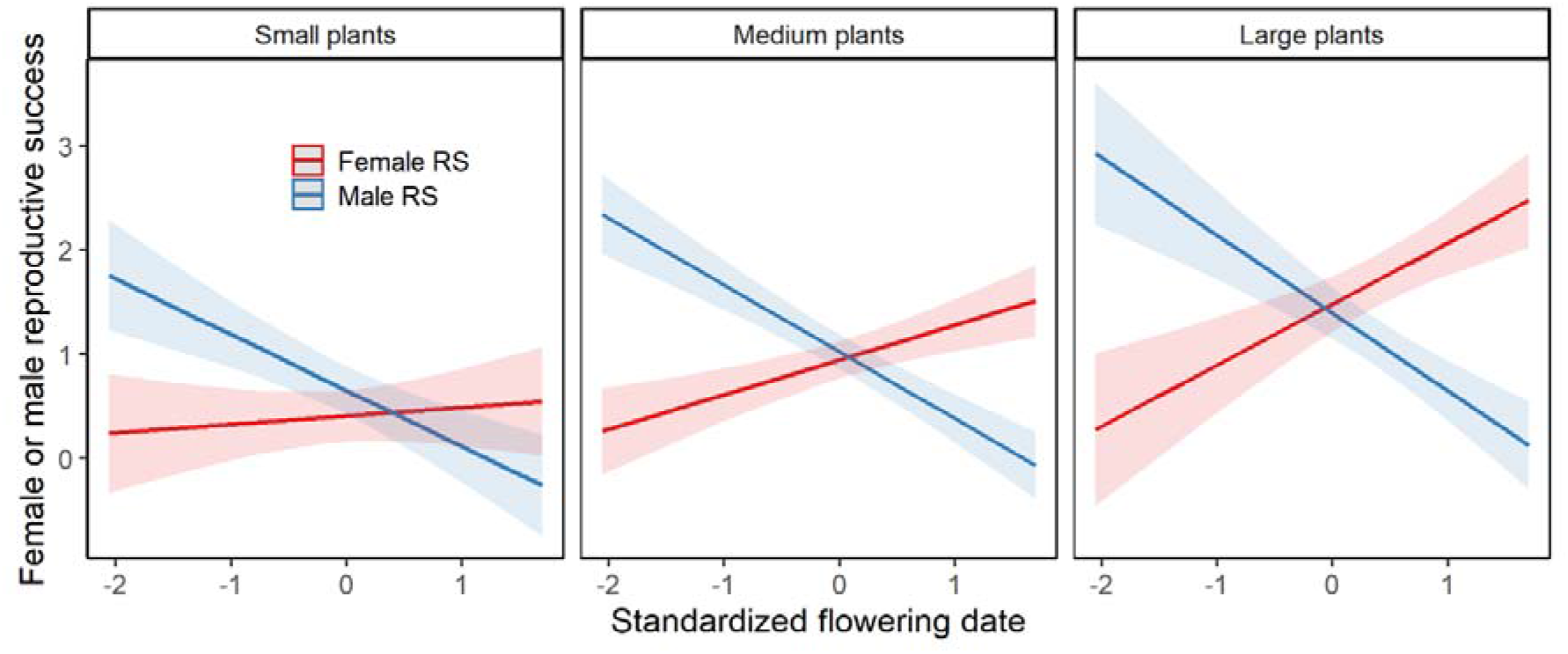
Plots showing the three-way interaction between size, flowering date, and sexual function on components of reproductive success (RS) in Population LM, as predicted by the model (see main text for details). Early flowering favors male RS for all plant sizes. However, female RS increases less steeply with flowering date in small plants (see Table S3; interaction among size, flowering date, and sex function: *P* < 0.05). The 95 % confidence interval of the estimates is shown around the regression lines.

## Notes

### Competing Interest Statement

The authors have declared no competing interest.

